# Engineering supramolecular organizing centers for optogenetic control of innate immune responses

**DOI:** 10.1101/2020.09.29.317776

**Authors:** Peng Tan, Lian He, Yubin Zhou

**Affiliations:** Institute of Biosciences and Technology, College of Medicine, Texas A&M University, Houston, TX 77030; Broad Institute of MIT and Harvard, Cambridge, MA 02142

**Keywords:** Synthetic biology, Optogenetics, Innate immunity

## Abstract

The spatiotemporal organization of oligomeric protein complexes and translocons, such as the supramolecular organizing centers (SMOC) made of MyDDosome and MAVSome, are essential for transcriptional activation of host inflammatory responses and immune metabolisms. Light-inducible assembly of MyDDosome and MAVSome are presented herein to induce activation of nuclear factor-kB (NF-κB) and type-I interferons (IFNs). Engineering of SMOCs and the downstream transcription factor permits programmable and customized innate immune operations in a light-dependent manner. These synthetic molecular tools will likely enable optical and user-defined modulation of innate immunity at a high spatiotemporal resolution to facilitate mechanistic studies of distinct modes of innate immune activations and potential intervention of immune disorders and cancer.

---

The optimal activation of innate immune responses for host defense requires the sensing of the invaded pathogen-associated molecular pattern (PAMPs), subcellular translocation of signaling molecules to trigger the assembly of large signaling aggregates or supramolecular organizing centers (SMOCs) (1), such as MAVSome, MyDDosome, and inflammasome. The activations of membrane toll-like receptors (TLRs) and cytosolic retinoic acid-inducible gene I (RIG-I)-like and nucleotide-binding and oligomerization domain (NOD)-like receptor signaling trigger the formation of SMOCs on various membrane-bound organelles (2) or other intracellular sites (3), SMOCs act as signaling platforms with locally concentrated signaling components to increase the response threshold for enzyme activation and assist signal amplification, which potentially defines the specificity of immune and metabolic responses (4). Activation of NF-κB and type I IFN pathways of MyDDosome and MAVSome leads to the nuclear translocation of the NF-κB complex and interferon regulatory factor 3/7 (IRF3/IRF7) to induce the expressions of tumor necrosis factor-α (TNF-α), interleukin-6 (IL-6) and IFNs. Subsequent binding of IFNs to IFNAR (5) and IL-6 to IL-6R-gp130 activate the STAT1/3 pathway (6) and the expression of IFN-stimulated genes and proinflammatory cytokines, ultimately mounting a robust immune response that could be beneficial for pathogen defense (7). Nevertheless, this host defensive reaction could also cause detrimental tissue injury or severe respiratory syndrome as seen in COVID-19 patients infected with SARS-CoV-2 (8). For example, IFNs stimulate the expression of SARS-CoV-2 entry receptor ACE2 (angiotensin-converting enzyme 2) to favor viral infection and IL-6-induced severe respiratory syndrome (9, 10). The association of ligand with TLR triggers the recruitment of TLR-specific Toll/IL-1R (TIR) domain-containing adaptor protein (TIRAP) on the plasma membrane (TLR4) and endosomal subdomains (TLR7/9) to induce MyD88 oligomerization (MyDDosome) through TIR domain interactions (11). MyD88 death domain (DD) and the intermediate domain (INT) are necessary for IL-1R-associated kinase (IRAK) binding, signaling transduction and downstream NF-κB and IRF7 activation (11, 12). Stimulation of different TLRs triggers the association of MyD88 and induces distinct patterns of gene expression (13, 14).

Recently, a study suggests that MyD88 could be rewired for glycolysis (4). Thus, customized MyD88 signaling could not only be used for the activation of innate immunity but also to instruct the development of antigen-specific acquired immunity and the acquired metabolism. Similarly, the binding of viral RNA to RIG-I triggers RIG-I K63 polyubiquitination and mitochondrial translocation to engage and induce MAVS (mitochondrial antiviral-signaling protein) prion-like polymers (MAVSome) (15), which in turn induces TANK binding kinase 1 (TBK1)-IRF3 interaction to trigger IFN production.(16) Currently, the studies of Toll-like and RIG-I-like receptor signaling use synthetic PAMP ligands, such as poly(I:C) for TLR3 and RIG-I, LPS for TLR4, and CpG for TLR9. However, these reagents lack high signaling specificity because they can activate multiple cross-talking signaling pathways. For instance, dsRNA activates IFN pathways involving TLR3 and RIG-I/MDA5, as well as PRK (protein kinase K), to cause translational suppression(17, 18), whereas dsDNA activates TLR9, AIM2 and cGAS (19). Potential ligand toxicity may also lead to acute cell death (20). There remains, therefore, a critical need for synthetic biology tools with minimal invasion to the host that could fine-tune the oligomerization and formation of MyDDosome and MAVSome. Such platforms will allow the precise control of the time, location and doses of innate immune responses to achieve tailored function.

Optogenetics, which incorporates synthetic photosensitive modules into cells of living tissues to probe biochemical pathways at specific subcellular locations with exquisite spatial and temporal control, provides an ideal solution to overcome the side effects associated with conventional TLR and RLR activating methods (21). Genetically encoded photo-switchable modules have been employed to control cellular signaling through modulating protein-protein interaction (e.g., LOVTRAP based on the ZDK1-LOV2 pair) (22), protein oligomerization (cryptochrome 2; CRY2) (23), and conformational switch (light-oxygen-voltage domain 2, LOV2) (24). For example, the LOVTRAP system has been exploited to control the subcellular release of target proteins from a particular subcellular organelle (e.g., mitochondria (25) via fusion with a mitochondrial anchoring sequence from Tom20) to control small GTPase activity and modulate cell edge protrusion (22). CRY2-or LOV2-fused Ca2+ channel actuators have been developed to engage ORAI1 channels to induce Ca2+ influx and NFAT signaling in cells of the immune system and tissue morphogenesis (21, 26–30). Inspired by these engineering strategies, we have developed a light-stimulable Photo-SMOC (MyDDosome and MAVSome) system and a light-controllable IRF3 activation device to photo-tune both the upstream and downstream signals that drive innate immune responses and cytokine production (Figure 1). Recent studies have shown that type I interferon (IFN) production by host antigen-presenting cells (APCs) and tumor cells can improve tumor immunogenicity and the sensitivity to immune checkpoint blockade in vivo (31, 32). Our optogenetic engineering methods will likely enable synthetic and programmable immune responses and set the stage for the development of optimized cancer immunotherapy, as well as the development of all-optical screening platforms for future identification of antimicrobial and immunomodulatory agents.

**Figure 1.**
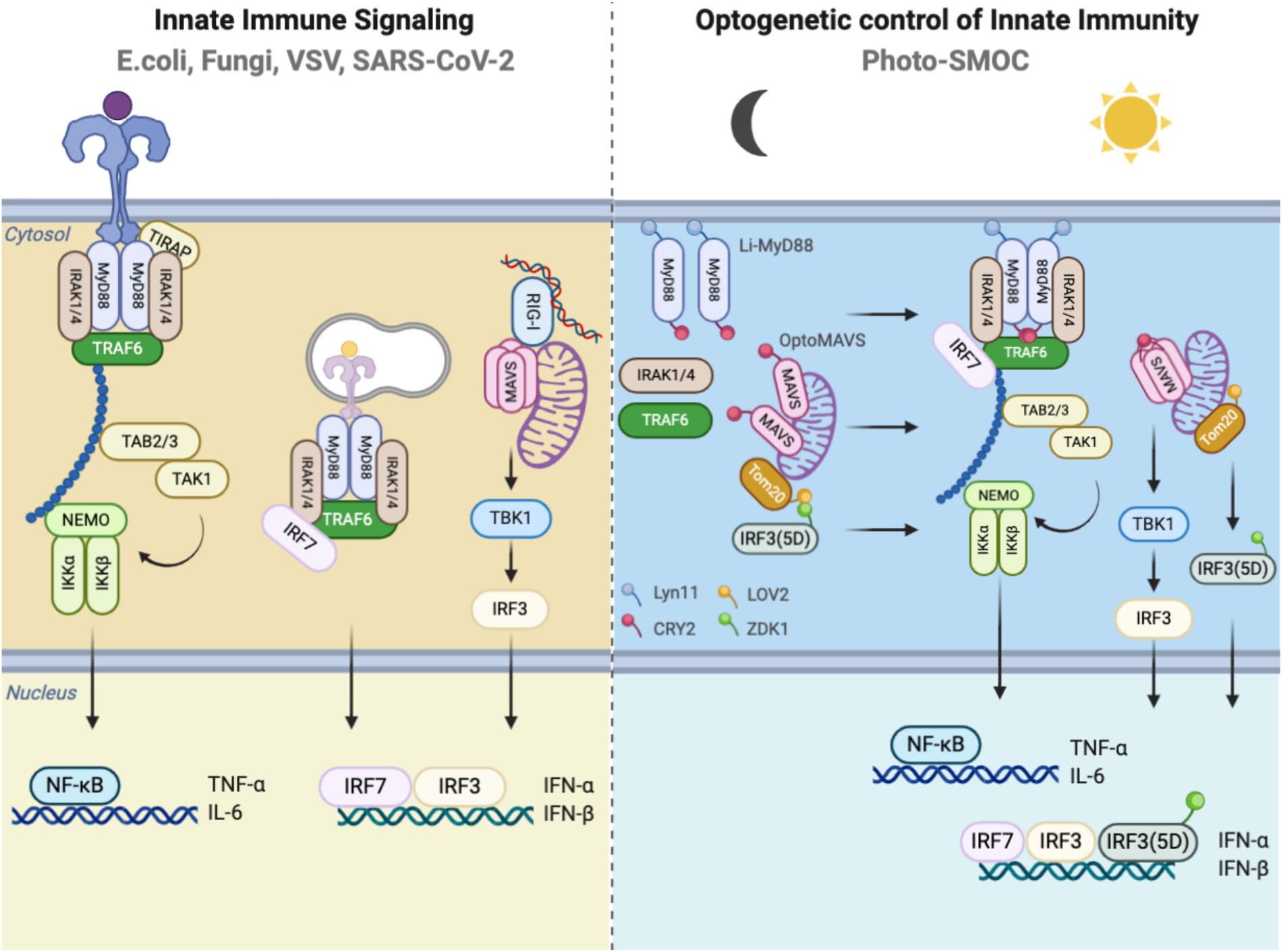
Photo-inducible SMOC (Photo-SMOC) assembly and IRF3 nuclear entry to photo-control innate immune responses.

## Li-MyD88 for photoactivatable boosting of TLR signaling

Optogenetics has recently been increasingly applied to remotely modulate biological processes other than the central nervous system, including the enteric nervous system (33) cancer immunotherapy (26, 34) ion channels (35, 36), interorganellar communication (37, 38) transcriptional reprogramming (39) and the CRISPR-based genome engineering (40), antibody and protein function (41). Recent efforts have been focusing on the engineering of controllable innate immune sensing and activation by opto- or chemo-genetics toward the goal of achieving customized immunomodulation (designated synthetic immunology) at high spatiotemporal resolution (42, 43). MyD88, as the most universal TLR adaptor, was selected as the prime candidate for generating optogenetic tools to modulate TLR activation.

To replace potentially toxic TLR ligands, we set out to design a single component optogenetic module that allows specific control of MyD88 oligomerization by taking advantage of the N-terminal photolyase homology region (PHR) of CRY2 from Arabidopsis thaliana. CRY2 is known to undergo light-dependent oligomerization within seconds in a reversible manner (44). Oligomerization of MyD88 leads to the recruitment and activation of the IRAK family of kinases via DD (death domain) interactions (45). Both the DD and INT (intermediary) domains, but not the TIR (Toll/Interleukin-1 receptor) domain, are required to trigger a signaling cascade that culminates in NF-κB activation (45, 46). The removal of he TIR domain is anticipated to reduce cross-talks with other signaling pathways. For instance, the TIR domain is required for the interaction of MyD88 with TLR and IL-1R receptors, and it also causes immune suppression when binding to microbial TIR domain-containing proteins (e.g. TIR hijacked by Bacterial TcpB for TLR suppression)(47), thereby acting as a limiting step for TLR/ NF-κB activation. Based on these facts, we fused CRY2 with a MyD88 fragment lacking the TIR domain (MyD88ΔTIR), and further appended a sequence motif derived from Lyn11 to facilitate its localization and oligomerization at the plasma membrane (PM) (48). In our envisioned design, we anticipated a robust blue light-induced multimerization of the synthetic protein (designated Li-MyD88; Lyn11-MyD88ΔTIR-CRY2-mCh) in the PM and subsequent boosting of the downstream activation of NF-κB and IRF7, as well as cytokine production (Figure 2A). Indeed, when expressed in HeLa cells, we observed a pronounced light-dependent clustering of MyD88 in the PM (Figure 2B), a process that could be readily reversed upon withdrawal of light stimulation (t1/2, activation = 1.6 ± 0.1 min; t1/2, deactivation = 5.3 ± 0.2 min; Figure 2C). To validate the light-induced consequences in mammalian cells, we transfected HeLa cells and HEK293T cells with a Li-MyD88 construct and measured the IKK phosphorylation with immunoblotting and the NF-κB promoter luciferase activity (Figure 2D-E). A basal level of IKK activation and NF-κB luciferase activity was observed (Figure 2D, lane 1 vs 3), likely due to plasmid transfection and MyD88 overexpression (49). Biochemically, we noted an appreciable light-dependent increase of the phosphorylation of IKK, which signals the activation of NF-κB signaling (Figure 2D). Functionally, we detected a 2.5-fold increase in the NF-κB luciferase activity in cells exposed to light stimulation (Figure 2D). We further moved on to test the light-induced effect on IRF7. It is known that endosomal TLRs activation results in the induction of IRF7 nucleus translocation and type I IFNs (IFN-α/β) in a MyD88-dependent manner in both HeLa and plasmacytoid dendritic cells (pDC) upon CpG ligand stimulation (12, 50). Interestingly, we observed endosomal TLR activation-independent but MyD88 oligomerization-dependent IRF7 nuclear translocation induced by light when Li-MyD88 and GFP-IRF7 were co-expressed in HeLa cells (Figure 2F). These results suggest that the oligomerization of MyD88, with the ensuing MyD88-IRF7 interaction, is sufficient to induce IRF7 nuclear shuttling for IFN induction. The activation of innate immune responses often shows high cell-type specificity and various amplitudes in different cells. For instance, TLR9 ligand A/D-type CpG oligodeoxynucleotide (CpG-A) can only activate the MyD88-IRF7 signaling axis in pDC but not in conventional DC (50). Our engineered Photo-SMOC tool provides a more generally-applicable approach to replace toxic ligands or stimulants while overcoming the strict requirement and cell-type-specific mechanisms to activate immune signaling (e.g. MyD88-IRF7 signaling). This desirable feature could be beneficial for drug candidate screening and regulator discovery without apparent toxicity and damage to the host cells.

**Figure 2.**
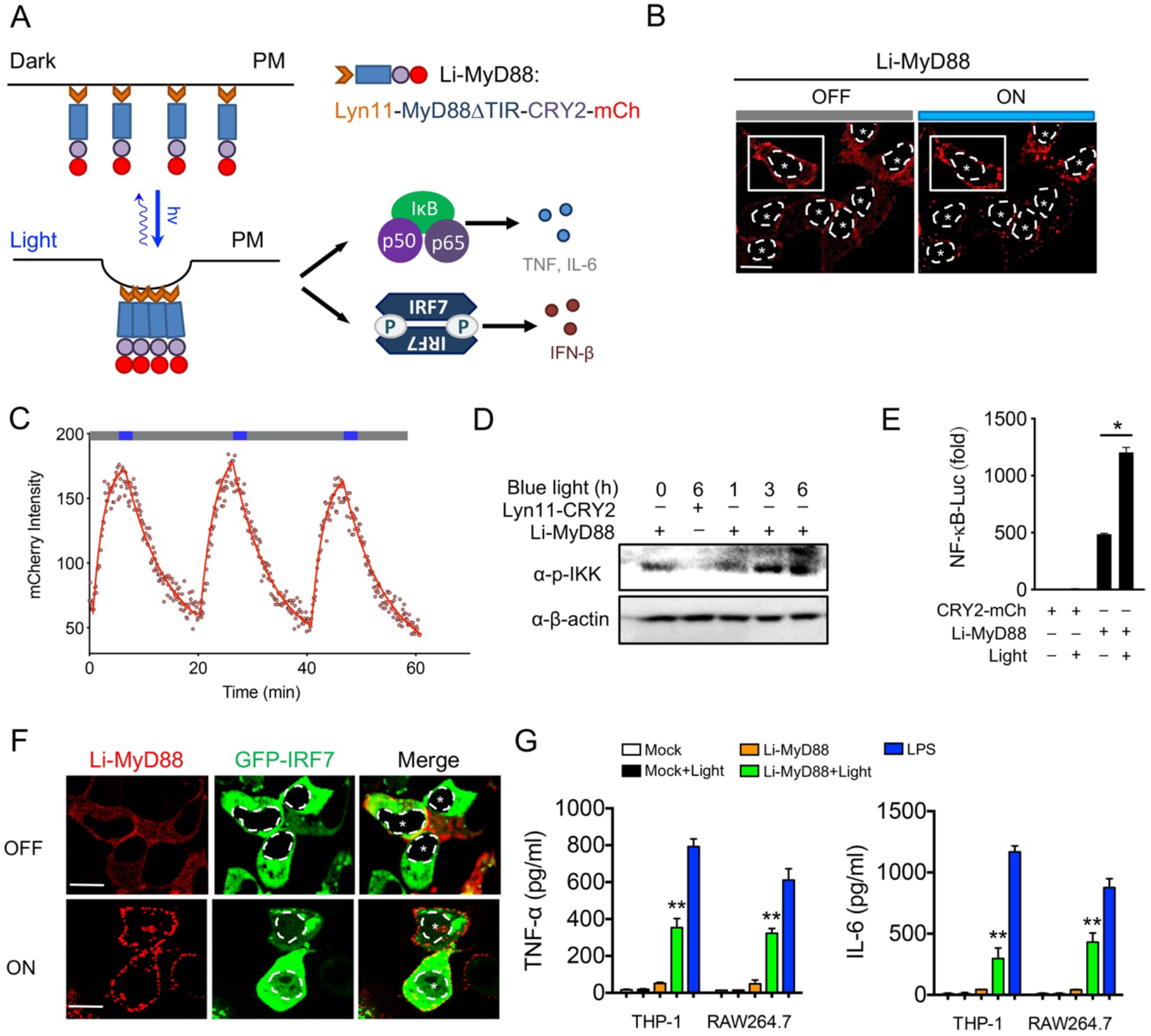
Design of a PM-tethered light-activatable MyD88 (Li-MyD88). Photostimulation was applied at 470 nm with a power density of 40 μW/mm^2^. Data were shown as means ± SD of at least three independent experiments. * *P* < 0.05, ** *P* < 0.01 (two-tailed paired Student’s *t*-test). A)Schematic of NF-κB and IRF7 activation and cytokine production through light-inducible oligomerization of PM-anchored MyD88 (designated Li-MyD88). B) Representative confocal images of HeLa cells showing Li-MyD88 puncta formation upon light illumination. Cells were exposed to blue light for 30 s. White dashed line and asterisks indicate the location of the nucleus. Signals in the boxed area were used for the quantification of Li-MyD88 clustering (shown in panel C). Scale bar, 20 μm. C) Quantification of reversible clustering of Li-MyD88 in the plasma membrane of HeLa cells. mCherry signals from a representative cell in (panel B, white box) were monitored over three repeated light-dark cycles (blue bar, ON; dark bar, OFF). D) Immunoblot analysis of IKK phosphorylation (p-IKK) in HeLa cells transfected with Lyn11-CRY2-mCh (control) or Li-MyD88. HeLa cells were exposed to pulsed blue light stimulation for 0, 1, 3, 6 h (470 nm, 30 s ON with 30 s interval; 40 μW/mm^2^). E) NF-κB promoter luciferase reporter assay of HEK293T cells transfected with CRY2-mCh (control) or Li-MyD88 with or without pulsed blue light illumination for 8 h. F) Representative confocal images of HeLa cells showing Li-MyD88 puncta formation (red) and nuclear translocation of GFP-IRF7 (green; the nuclear envelope demarcated by a dashed line) upon blue light illumination. Cells were exposed to pulsed blue light for 2 h. Scale bar, 20 μm. G) Measurement of TNF-α and IL-6 production by ELISA in the supernatants of THP-1 and RAW264.7 cells expressing an empty vector (control) or Li-MyD88 with or without pulsed light illumination for 12 h. 100 ng/ml LPS stimulation for 12 h was used as a positive control.

To further test the functional consequence of Li-MyD88 in more physiologically-relevant contexts, we transfected human monocyte THP1 cells and mouse macrophage RAW264.7 cells with Li-MyD88 or the empty vector as control. TNF-α and IL-6 production in these two cell types, before and after photostimulation, was assessed as a readout for the activation of NF-κB singling. We found that Li-MyD88 was able to induce both TNF-α and IL-6 production upon blue light (Figure 2G), but to a lesser extent when compared with LPS stimulation. This discrepancy might be attributed to the varying doses of light and LPS, as well as the difference in the transfection efficiency of the Li-MyD88 construct. Collectively, these results establish Li-MyD88 as a genetically-encoded innate immunity modulator that allows photoactivatable boosting of the TLR-mediated signaling pathway.

## Light-inducible assembly of MAVS signalosome for IFN activation

Viral infection induces the RIG-I activation, which in turn catalyzes the conversion of MAVS into prion-like aggregates on the mitochondrial membrane in the presence of K63 ubiquitin chains (15). Earlier studies have shown that overexpression of any of the key signaling components in the RIG-I-MAVS signaling pathway could induce IRF3 activation and IFN production, suggesting concentration-dependent signalosome self-assembly(2, 51). To test whether MAVS alone could form functional polymers via in situ manipulation of MAVS oligomerization in real-time, we decided to generate a chimeric construct by fusing CRY2 with MAVS (OptoMAVS; Figure 3A). Upon blue light stimulation, mitochondrial MAVS fibers started to concentrate and cluster, a process that could be reversed after switching off the light source (Figure 3B). We next tested if OptoMAVS could activate the downstream IRF3 activation as evaluated by its phosphorylation with immunoblotting. Blue light stimulation indeed triggered a marked increase of IRF3 phosphorylation in HeLa cells expressing optoMAVS but not in those expressing the wild-type MAVS (Figure 3C). More importantly, OptoMAVS-expressing cells showed a light-dependent boosting (>10-fold) of the interferon-stimulated response element (ISRE)- and IFN-β luciferase activity (Figure 3D), two commonly used reporters to examine IFN activation (52). In THP-1 and RAW cells, OptoMAVS induced efficient IFN-β production after light stimulation, at a level that was comparable to VSV stimulation (Figure 3E). Together, our results demonstrate the generation of photo-inducible MAVS-derived SMOC that is capable of photo-triggering type I IFN activation.

**Figure 3.**
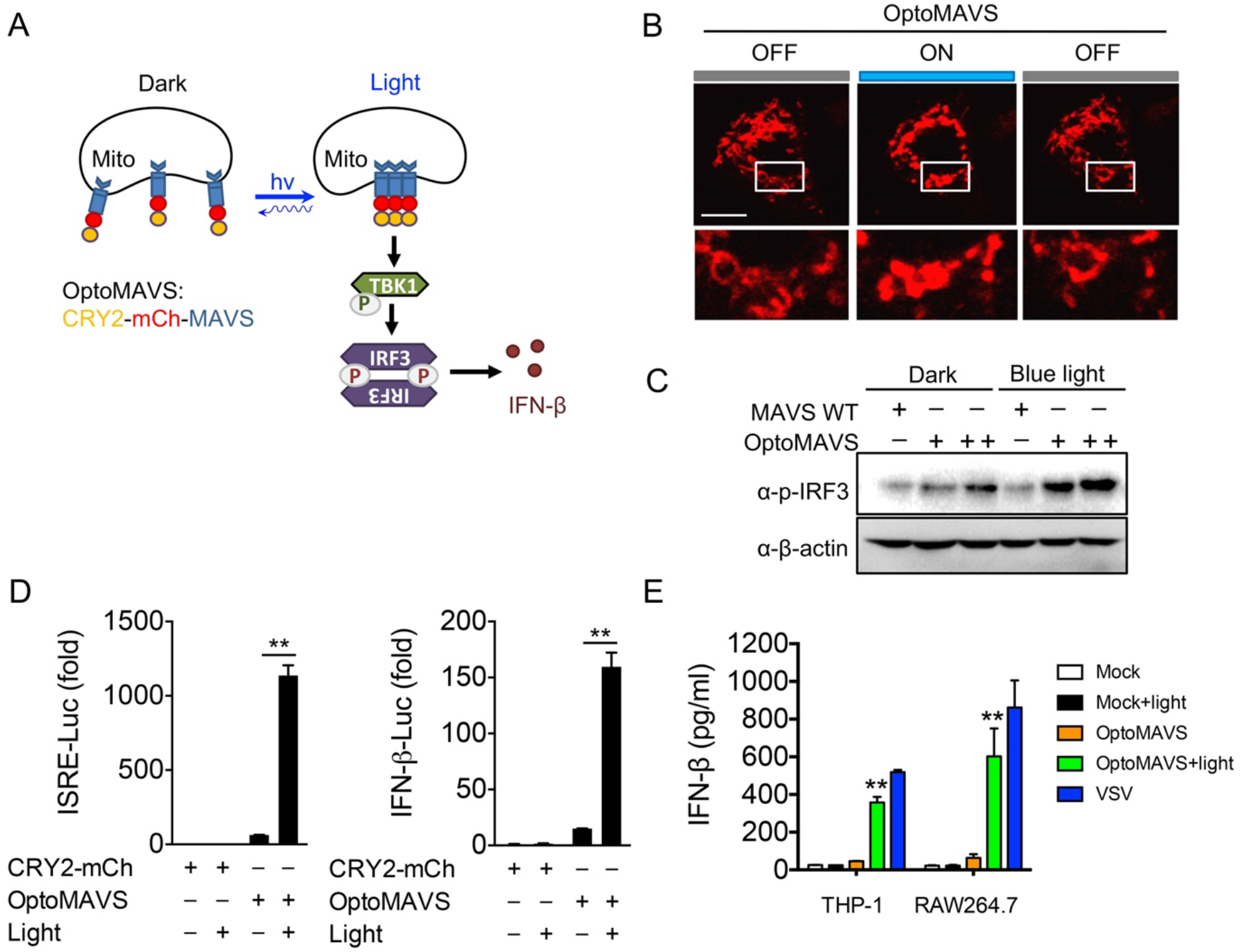
Design of a light-sensitive mitochondrial MAVS signalosome. Photostimulation was applied at 470 nm with a power density of 40 μW/mm^2^. Data were shown as means ± SD (n=3). ** *P* < 0.01 (two-tailed paired Student’s *t*-test). A) Schematic of IRF3 activation by TBK1 through light-inducible mitochondrial MAVS oligomerization (designated OptoMAVS; CRY2-mCh-MAVS). B) Confocal imaging showing reversible, light-inducible clustering of mitochondrial OptoMAVS in HeLa cells. Inset, zoomed-in views of the boxed areas. Photostimulation was applied at 470 nm (40 μW/mm^2^, ON for 10 s, OFF for 5 min). Scale Bar, 20 μm. C) Immunoblot analysis of IRF3 phosphorylation (p-IRF3) in HeLa cells transfected with WT MAVS or CRY2-MAVS (+, 20 ng) or CRY2-MAVS (++, 80 ng) with pulsed blue light for 8 h. D) ISRE and IFN-β promoter-driven luciferase reporter assay in HEK293T cells transfected with CRY2-mCh (control) or OptoMAVS with or without pulsed blue light irradiation for 8 h. E) ELISA measurement of IFN-β production in the supernatants of THP-1 and RAW264.7 cells expressing the empty vector (control) or OptoMAVS with or without pulsed light illumination for 12 h. VSV (MOI = 1) stimulation for 12 h was used as a positive control.

Recent studies highlight the diverse functional outputs of SMOC signaling beyond the immune function and hint the possibility to create unique signaling circuits (4). For example, TBK1 serves as a component of MyDDosome in TLR-dependent glycolysis, and therefore, MyD88 could be rewired to induce TBK1 activation and IFN response by inserting a pLxIS motif embedded within MAVS, TRIF and STING (16). It is likely that our Photo-SMOCs could be used to identify key components in SMOC through monitoring the co-clustering of candidate proteins with MyD88 or MAVS. Moreover, the organization of SMOC often occurs on the specific cellular membranes, suggesting the evolution of SMOC to gain a specified function. Thus, it would be interesting to investigate the physiological output of Photo-SMOC when anchored to distinct subcellular compartments in follow-on studies.

## Photo-controllable nucleocytoplasmic shuttling of IRF3 to modulate IFN transcription

A constitutively active form of IRF3(5D) (S396D, S398D, S402D, T404D, and S405D) has been shown to accumulate within the nuclei to drive the activation of IFN (53), which has been used as a screening platform for identifying SARS-CoV viral proteases for immune escape and drug development (54). Capitalizing on a LOVTRAP system (made of the LOV2-ZDK1 pair) that can induce protein-protein dissociation upon photostimulation (22), we sought to constrain IRF3(5D) toward the mitochondrial outer membrane in the dark and then release it following light stimulation to cause nuclear translocation with subsequent IFN activation (Figure 4A). Although the LOV2 domain has been used to cage the activity of the constitutive active form of a transcription factor or proteins with nuclear localization signals (NLS) to achieve light-induced nuclear translocation (55), it often requires the fusion of the effector domain to the C-terminus of LOV2. This class of optogenetic tools also tends to cause leaky nuclear localization due to incomplete caging of the effector domain or the NLS, a process also known as “dark activity”. As a result, intensive optimization steps (e.g., laborious optimization of the linker and mutagenesis of the interface residues between the LOV2 and the effector domain) are required to fully sequester the effector function. By contrast, the modular LOVTRAP system could be conveniently used for the rapid translocation of target proteins with minimized dark activity due to its tight interaction between LOV2 and ZDK1 in the dark and the high dynamic changes between the dark and lit states. The dark activity could thus be greatly mitigated without the need for extensive optogenetic engineering (56). Using this strategy, we tested if ZDK1-IRF3(5D) could be trapped nearby the mitochondrial outer membrane in the dark and then released to the cytosol upon light stimulation. As expected, we found that ZDK1-IRF3(5D) stayed anchored to the mitochondrial membrane as reflected in a tight colocalization with Tom20-Venus-LOV2. Following light stimulation, the majority of the mitochondria-anchored IRF3(5D) was released and dispersed into the cytosol (Figure 4B). This process could be readily reversed following the withdrawal of light, with the activation and deactivation kinetics determined to be 5.8 ± 0.8 s and 45.2 ± 4.6 s, respectively. With prolonged light stimulation, we also observed a gradual accumulation of IRF3(5D) in the nuclei within 2 hours (Figure 4C). Next, we used an IFN-β promoter-luciferase reporter assay to assess the light-inducible activation of IRF3-dependent IFN transcription. As expected, we noted a blue light-dependent activation of reporter gene expression in cells co-expressing ZDK1-IRF3(5D) and mitochondria-tethered LOV2 (Figure 4D). Taken together, our data suggest a reversible control of the cytosol-to-nucleus shuttling of IRF3 and inducible IFN signaling activation by light.

**Figure 4.**
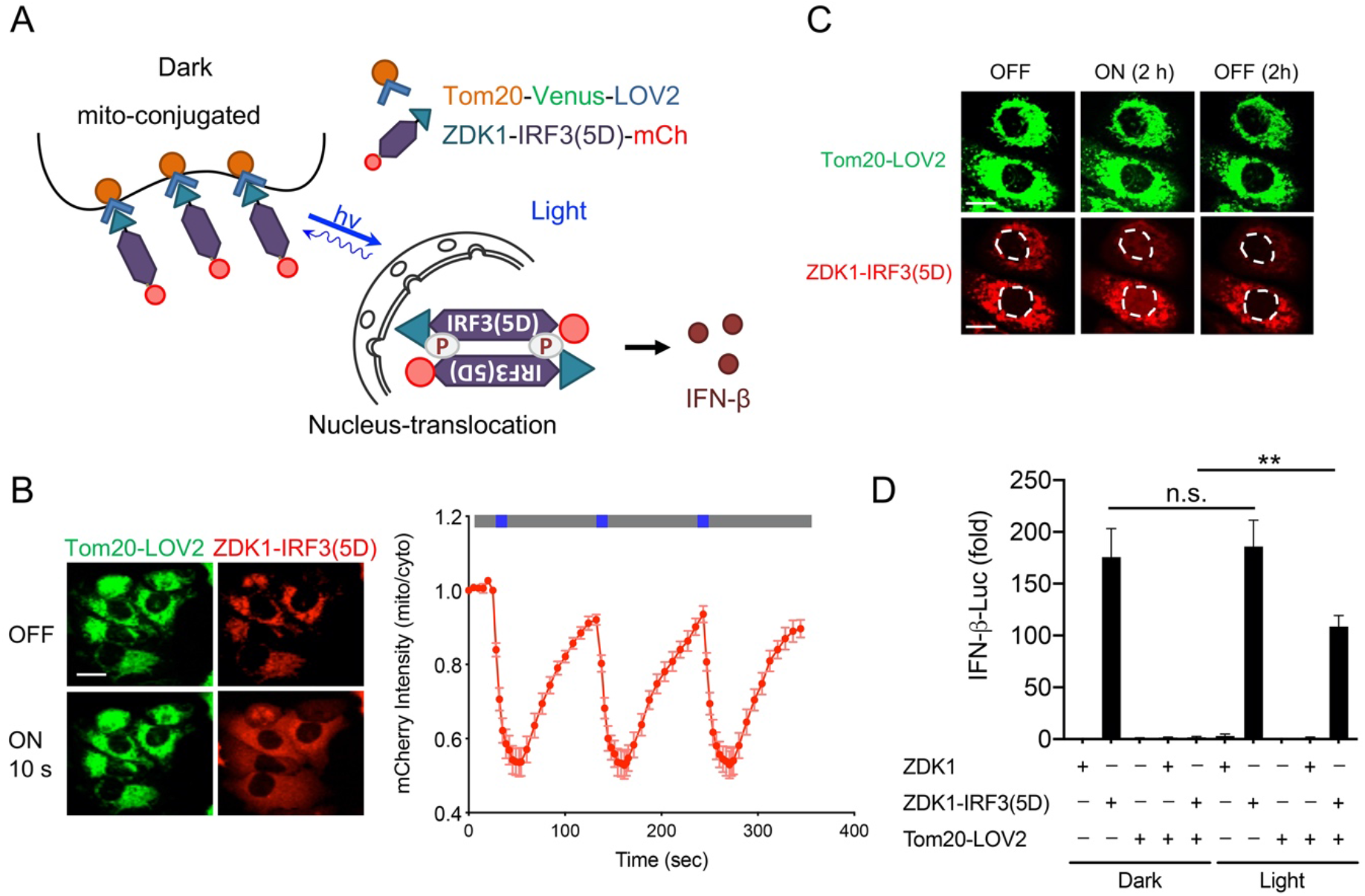
Light-inducible nuclear translocation of IRF3 to activate IFN-dependent gene transcription. Photostimulation was applied at 470 nm with a power density of 40 μW/mm^2^. A) Schematic of light-inducible release of ZDK1-IRF3(5D)-mCh from mitochondria-resident Tom20-Venus-LOV2 toward the cytosol and nucleus to activate IFN production. B) Confocal images of HeLa cells co-expressing Tom20-Venus-LOV2 (green) and ZDK1-IRF3(5D)-mCh (red) before and after exposure to blue light for 10 s. Scale bar, 20 μm. Quantification of mitochondrial mCherry signals over three repeated light-dark cycles (470 nm, 40 μW/mm^2^; blue bar, ON; dark bar, OFF). C) Representative confocal images of HeLa cells co-expressing Tom20-Venus-LOV2 and ZDK1-IRF3(5D)-mCh (red) with or without pulsed blue light illumination for 2 h. 2 h after switching off the light source, images were re-acquired to examine the reversibility of nuclear translocation. The nuclei were circled by white dashed lines. D) IFN-β promoter-luciferase reporter assay in HEK293T cells transfected with ZDK1-mCh or ZDK1-IRF3(5D)-mCh with or without co-expression of Tom20-Venus-LOV2. Transfected cells were either kept in the dark or subjected to pulsed blue light illumination for 12 h. Data were shown as means ± SD from three independent experiments. \*\**P* < 0.01, and n.s., not significant. (two-tailed paired Student’s *t*-test).

Photo-SMOC enables less toxic synthetic immunomodulation with improved specificity and spatial precision. Conventional immune activation by PAMP often lacks the signaling specificity because of molecular crosstalks among different innate sensing pathways. Multiple innate immune sensors can be activated by the same PAMP or DAMP. Meanwhile, multiple innate sensing pathways could involve overlapping signaling transducers and transcriptional programs (57). In addition, innate sensors are generally expressed in a wide variety of cell types, including immune cells, epithelial cells and cancer cells. These shared features by multiple signaling pathways and cell types pose challenges to signaling specificity and spatial precision, which are linked to on-target toxicity at non-diseased sites to cause systemic cytokine release syndrome and even cell death. To demonstrate the high signaling specificity, low cytotoxicity and spatiotemporal resolution of the Photo-SMOC system, we carried out a side-by-side comparison between light-induced Photo-SMOC activation (OptoMAVS as a test case) and synthetic PAMP stimulation. we transfected THP-1 cells with OptoMAVS (for photostimulation) or CRY2-mCh (as a control for PAMP stimulation), followed by sorting of mCh+ cells and PMA-priming. Synthetic poly(dA:dT) was used as a representative PAMP stimulus, which was found to not only activate the RIG-I/MAVS type I IFN pathway(58) but also the AIM2-ASC inflammasome pathway(59), thus leading to caspase-1 activation and subsequent cell death (Figure 4A-C) (59). By contrast, OptoMAVS specifically induced type I IFN signaling, as reported by the nuclear translocation of IRF3 (Figure 5A), without co-activating ASC inflammasome formation (Figure 5B). Compared to poly(dA:dT) that caused appreciable cell death as indicated by positive SYTOX staining (Figure 5C), we did not detect overt cytotoxicity in OptoMAVS-expressing cells after light stimulation (Figure 5C). Lastly, to demonstrate the spatial control of OptoMAVS, we locally illuminated a selected single cell while sparing other neighboring cells by using the FRAP module in a Nikon imaging system (488-nm; with the laser power output set at lower than 0.5%), followed by global light stimulation on the whole imaging field to activate all cells (Figure 5D). Our results showed that only the pre-stimulated single cell, but not the surrounding un-illuminated cells, showed IRF3 nuclear entry (Figure 5D, left). Subsequent global photo-illumination led to the nuclear translocation of IRF3 of cells under the imaging field (Figure 5D, right). Congruently, these results attest tot he high specificity, superior spatial precision and low cytotoxicity of the Phot-SMOC tool for synthetic immunomodulation. This feature will likely pave the way for precise manipulation of IFN-mediated transcriptional programs at defined tissue regions and specific cell types by harness the power of light.

**Figure 5.**
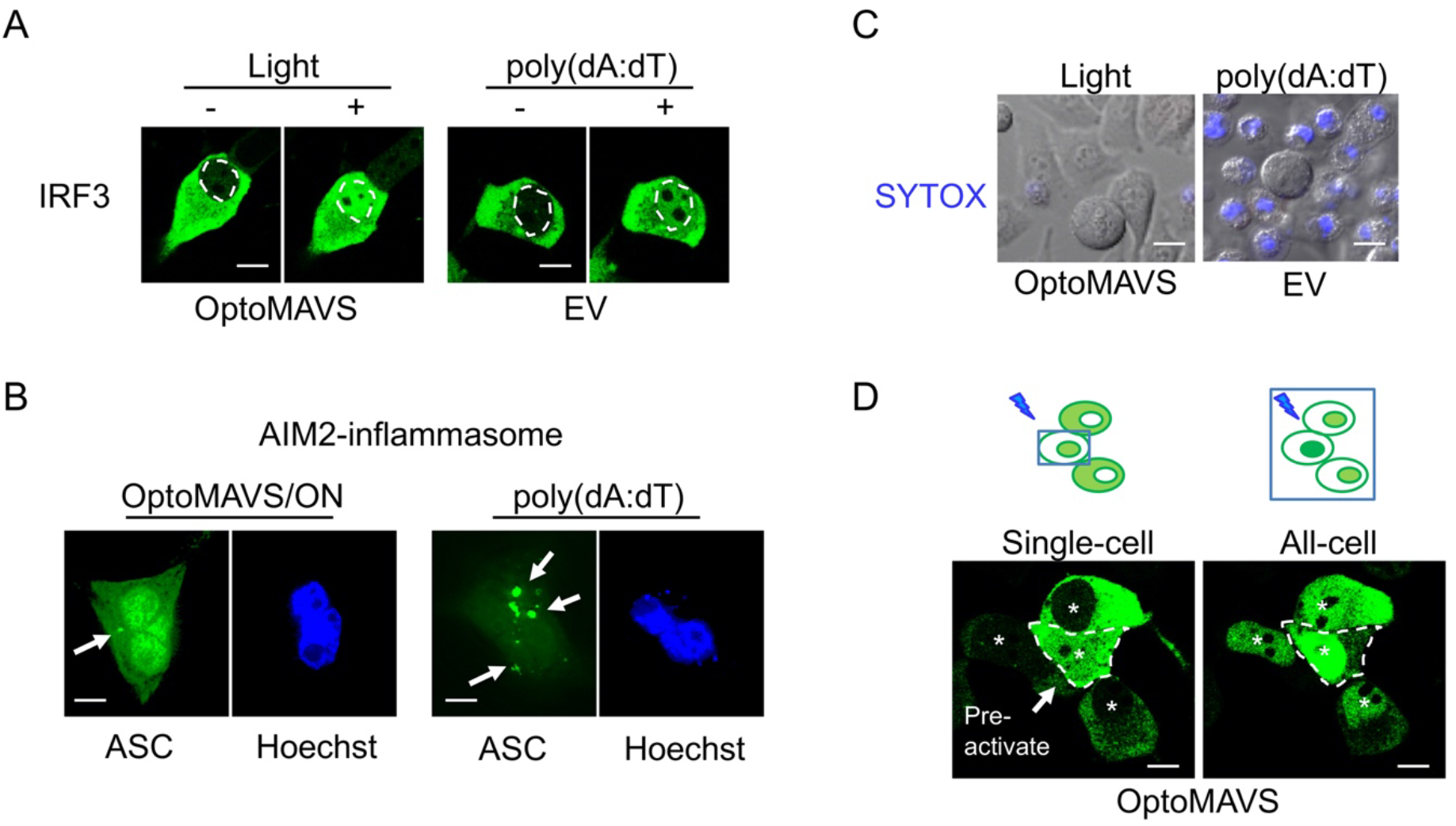
Comparing the specificity, cytotoxicity and spatial precision between OptoMAVS and conventional innate immune activation using poly(dA:dT). Photostimulation was applied at 470 nm with a mild power density of 40 μW/mm^2^. A) Scale bar, 10 μm. Representative confocal images showing the nuclear translocation of IRF3 (green) in PMA-primed (20 ng/mL) THP-1 cells transfected with OptoMAVS (for blue light pulse) or the CRY2-mCh empty vector (EV; 2μg/mL poly(dA:dT) stimulation for 6 h). The dashed line indicates the border between the cytosol and the nucleus. B) Representative confocal images showing ASC staining (green) in PMA-primed THP-1 cells. Cells were transfected with OptoMAVS (left) or CRY2-mCh (right), followed by blue light pulse (left) or 2μg/mL poly(dA:dT) stimulation (right) for 6 h. ACS speckles (indicated by arrows) were more pronouncedly detected in the poly(dA:dT) group. Nucleus (blue) was stained with Hoechst 33342. C) Assessment of cell death by SYTOX blue staining in PMA-primed THP-1 cells transfected with OptoMAVS or CRY2-mCh, followed by blue light pulse (left) or 2μg/mL poly(dA:dT) stimulation (right) for 6 h. D) Confocal images of spatially controlled IRF3 nuclear translocation. Photosimulation was first applied to a single cell (circled by a dashed line; left) for 3 h, followed by global light illumination upon the whole imaging field (right) for 3h. Asterisks indicate nuclei.

In this study, we engineered photoactivatable SMOCs (1) to control the activation of both TLR and RLR signaling pathways involved in innate immune responses. We provide solid evidence to support the notion that MyD88 and MAVS are the core units of SMOC, oligomerization of which alone is sufficient to activate the downstream inflammatory signaling and proinflammatory cytokine production in both human and mouse immune cells. We further adapted the LOVTRAP system (22) to modulate the intracellular localization of an essential transcription factor, IRF3, to induce innate type I IFN immune response. Our study expands the repertoire of molecular tools tailored for synthetic immune regulation (synthetic immunology) and a better understanding of the mechanisms underlying SMOC assembly. These tools will likely set the stage for future exploration of potential regulators and putative therapeutic avenues for treating immune-related diseases, as well as for exploring programmable features of other SMOCs within and beyond the innate immune systems, such as eliciting tumor inflammation to overcome cancer resistance toward immune checkpoint blockade therapy.

## Materials and Methods

### Cell culture and transfection

HeLa, HEK293T, THP-1, and RAW264.7 cells were obtained from American Type Culture Collection and cultured in Dulbecco’s modified Eagle’s media (DMEM, Sigma-Aldrich, St Louis, MO, USA) or RPMI-1640 medium, supplemented with 10% fetal bovine serum (FBS) under 37 °C at a 5% CO_2_ atmosphere. Phorbol 12-myristate 12-acetate (PMA) from Sigma Aldrich (St Louis, MO, USA) was used at 20 ng/mL to induce THP-1 differentiation to macrophage. DNA transfections were carried out by using the Lipofectamine 3000 reagent (Life Technologies, Carlsbad, CA, USA) or using an Amaxa nucleofector kit (Lonza) for transfection of plasmids into THP-1 and RAW264.7 cells. For live-cell imaging, cells were seeded in four-chamber 35-mm glass-bottom dishes (D35C4-20-1.5-N, Cellvis, Mountain View, CA, USA) at 40-60% confluency, and imaged 24 hours after transfection.

### Luciferase reporter assay

HEK293T (2×10^5^) cells were seeded into 24-well plates the day before transfection. Cells were co-transfected with the NF-κB, IFN-β, and ISRE promoter luciferase reporter and *Renilla* luciferase as a internal control (pRL-TK) using Lipofectamine 3000 (Invitrogen), along with Li-MyD88, OptoMAVS, Tom20-Venus-LOV2, ZDK1-IRF3(5D)-mCh (with Lyn11-CRY2-mCh, CRY2-mCh, ZDK1-IRF3(WT)-mCh as control) for reporter luciferase assay. 12 h after transfection, an external blue LED light source (470 nm, 40 μW/mm^2^) was used with the photo-cycles set as follows: ON for 30 sec on and OFF for 100 sec over the duration of 8 h. The duration and frequency of light pulses were controlled from an Arduino Uno board to the LED current driver.

Luciferase activity was determined using the Dual-Luciferase Assay (Promega) with the Luminoskan Ascent Luminometer (Thermo Scientific) as previously described (*2*). Reporter gene activity was normalized to the internal control.

### Assays for cytokine production

TNF-α, IL-6, and IFN-β were detected using an ELISA kit (Thermo Scientific) as previously described (*26*). Briefly, diluted supernatants and standards were applied to the pre-coated 96-well plate and incubated for 2 h at room temperature (RT) or overnight at 4 °C. The plate was then washed and incubated with biotin-conjugated detection antibody (1:1000) for 1 h at room temperature. Next, the plate was washed and incubated with HRP conjugate concentrate for 30 min. The plate was washed and incubated with the tetramethylbenzidine (TMB) substrate solution (Sigma-Aldrich). The reaction was stopped with 2 M H_2_SO4. The absorbance of each well was recorded at 450 nm. The absorbance of the standard sample was used to generate the standard curve.

### Immunoblot analysis

HEK293T or HeLa cells transfected with optogenetic constructs (dark and light) were lysed in low salt lysis buffer (50 mM Tris, pH7.5; 150 mM NaCl; 1% Triton-X; 5 mM EDTA; 10% (v/v) glycerol with protease inhibitor cocktail [Roche]) or RIPA buffer (150 mM NaCl, 1.0% IGEPAL CA-630, 0.5% sodium deoxycholate, 0.1% SDS, 50 mM Tris, pH 8.0 protease inhibitor cocktail [Roche]) and shook on ice for 15 min. Whole-cell lysates were boiled with 4x SDS loading buffer, and subjected to SDS-PAGE (Bio-Rad). Proteins were transferred to nitrocellulose membranes (Bio-Rad). Membranes were blocked in 5% dry milk/TBST for 1 h at RT and incubated with p-IKK, p-IRF3, mCherry or β-actin antibodies (Cell Signaling Technology). Membranes were developed using the Luminata Western HRP Chemiluminescence Substrates (Millipore) and ChemiDoc XRS+ System with Image Lab (Bio-rad).

### Immunofluorescence assay and live-cell imaging

HeLa cells were either left untreated or treated with blue light followed by fixation in 4% paraformaldehyde solution in PBS at RT for 15 min and permeabilized at room temperature with 0.1% Triton X-100/PBS. The Nikon A1R confocal module mounted onto an inverted Nikon Eclipse Ti-E body was used for confocal imaging. Nikon multi-line argon laser sources (405/488/561/640 nm) were used to excite the corresponding fluorophores. The 40x and 60x objectives (Nikon) were used for image acquisition. All acquired images were analyzed and the correlation coefficient (r) of pixel intensity values was extracted by using the Nikon NIS-Elements AR package or the ImageJ (NIH) software. For live-cell imaging, a caged platform was used to maintain the proper temperature, CO_2_ and humidity to keep cells healthy during the imaging process. To monitor repeated oligomerization of Li-MyD88 and OptoMAVS, the association and dissociation of Tom20-Venus-LOV2/ZDK1-IRF3(5D)-mCh, confocal images of HeLa co-transfected with these optogenetic plasmids were acquired using the 488-nm laser source to excite GFP (with 0.1-5% output). To monitor ZDK1-IRF3(5D)-mCh and IRF7-GFP nucleus translocation, an external blue LED light source (470 nm, 40 μW/mm^2^) was used with the photo-cycles set as follows: ON for 30 sec on and OFF for 30 sec over the duration of 2 h.

### Generation of optogenetic constructs

For the generation of Li-MyD88 plasmid, the cDNA sequence encoding MyD88ΔTIR was PCR amplified using primers (BamHI-MyD88 ΔTIR-FWD: 5’-CGCGGATCCCGACCCGACCGCGCTGAG-3’; EcoRI-MyD88 ΔTIR-REV: 5’ CCGGAATTCCGCCAGCTCTGCTGTCCG-3’) and inserted to the 3’ end of pcDNA3.1-Lyn11 using BamHI and EcoRI. Lyn11-MyD88 ΔTIR fragment was amplified by PCR flanking with two NheI enzyme cutting sites using primers (NheI-Lyn11-FWD: 5’-CTAGCTAGCATGGGATGTATAAAATC-3’; NheI-MyD88 ΔTIR-REV: 5’-CTAGCTAGCTTGAATTCCGCCAGCTCTGCTGTCCG-3’) and inserted to the NheI site of pmCherry2-CRY2 to get pmCherry-Lyn11-MyD88 ΔTIR-CRY2 (Li-MyD88). Correct insertion was confirmed by sanger sequencing.

To generate OptoMAVS plasmid, the cDNA sequences encoding full-length MAVS was amplified from pENTR-MAVS with GS linker (GGSGG) at 5’ end by PCR flanking with two SmaI sites using primers (CRY2-hMAVS-mCherry-FWD: 5’-TCCCCCGGGGCGGATCTGGTGGCCCGTTTGCTGAAGACAAGACC-3’; CRY2-hMAVS-mCherry-REV: 5’-TCCCCCGGGAGTGCAGACGCCGCCGGTACAG-3’) and inserted to SmaI site of pmCherry2-CRY2 to get pmCherry-MAVS-CRY2 (OptoMAVS). To generate ZDK1-IRF3(WT/5D)-mCh plasmid, the cDNA sequences encoding IRF3(WT) or IRF3(5D) was amplified from pcDNA3-IRF3(WT) or pcDNA3-IRF3(5D) by PCR flanking with two NotI enzyme cutting sites using primers (NotI-IRF3(5D)-FWD: 5’-ATAAGAATGCGGCCGCATGGGAACCCCAAAGCCAC-3’; NotI-IRF3(5D)-REV: 5’-ATAAGAATGCGGCCGCTCAGCTCTCCCCAGGGCCC-3’) and inserted to NotI site of pTriEX-ZDK1-mCh-6His to get pTriEX-ZDK1-mCh-IRF3(5D). Correct insertion was confirmed by sanger sequencing.

### Imaging data analysis

Images were analyzed with the Nikon NIS-Elements imaging software. To quantify the fluorescent signals of photoactivated oligomerization and protein translocation at selected areas, we used the region-of-interest (ROI) toolbox in Nikon NIS-Elements software to define the plasma membrane, cytosolic, mitochondrial and nucleus regions manually with 20-30 cells selected for each analysis. The fluorescence intensity ratio (F_mito_/F_cyto_) was used as a readout for assessing mitochondria-to-cytosol translocation. The “Time Measurement” tool was used to determine the mCherry intensities for Li-MyD88 oligomerization and LOVTRAP IRF3(5D) relocation. For spatially-restricted photostimulation, the FRAP module in the Nikon imaging system was used, with the 488-nm laser power output set at lower than 0.5%.

### Statistical analysis

Data were analyzed by two-tailed Student’s t-test or ANOVA for multiple comparisons. Data were presented as mean ± standard error (SD). The sample size for each experiment, n, was included in the results section and the associated figure legend. All analyses were performed with GraphPad Prism 5 (GraphPad Software, La Jolla, CA). *P* values <0.05 were considered significant.

## Acknowledgments

We thank Dr. Douglas Golenbock at UMass Medical School for sharing GFP-tagged IRFs constructs, Dr. Klaus Hahn for sharing the LOVTRAP system. This work was supported by the Welch Foundation (BE-1913-20190330 to Y.Z.), the American Cancer Society (RSG-16-215-01-TBE to Y.Z.), the National Institutes of Health (R01GM112003 to Y.Z., R21GM126532, and R01CA232017 to Y.Z.), and the John S. Dunn Foundation (to Y.Z.).

## Author Contributions

P.T. and Y.Z. conceived the project. P.T. and L.H. constructed optogenetic constructs and performed imaging and data analysis. Y.Z. supervised the entire project. P.T. and Y.Z. wrote the manuscript. All authors edited the manuscript.

